# MEG-informed navigated TMS for individualized speech cortical mapping

**DOI:** 10.64898/2026.07.10.737657

**Authors:** Salla Autti, Sara Korkealaakso, Juha Gogulski, Melina Engelhardt, Selja Vaalto, Hanna Renvall, Mia Liljeström, Pantelis Lioumis

## Abstract

Speech cortical mapping by means of navigated repetitive transcranial magnetic stimulation (SCM nrTMS) provides neurosurgeons with noninvasive prior information about individual’s cortical speech network. Individualized mapping is required, since the exact locations and activation patterns of speech production show high variability between individuals. We hypothesized that magnetoencephalography (MEG) data of an individual’s speech production could guide the SCM TMS process temporally and spatially, leading to higher error rates at MEG-defined locations with TMS pulse timings coinciding with MEG activity. 13 healthy subjects participated in MEG and TMS measurements, where the timing of the TMS pulse (PTI; picture-to-TMS interval) was adjusted based on the individual’s MEG activation in a picture naming task. At the group level, significant correlations were observed between the latency of the peak MEG activation and the PTI that produced the highest speech error rate. The MEG peak preceded the best PTI by 132 ms (R=0.713, *p*=0.006) across the entire stimulation area in the lateral left hemisphere, and by 103 ms (R=0.673, *p*=0.012) in the left frontal regions. We found 17 combinations of PTI and stimulation area in which the subject’s speech error rate increased significantly compared to their average error rate. Our findings suggest that optimal PTIs are highly individual, and that individualizing the PTI according to MEG activation provides a straightforward method for accounting individual variability in speech function and may increase the sensitivity and utility of SCM TMS.

## Introduction

During neurosurgery, the extent of tissue resection must be optimized to preserve brain regions that support critical functions for patient integrity and quality of life. Accurate identification of language-related brain regions prior to resection is of the highest priority but challenging, as functional areas cannot be identified at the individual level based on mere anatomical landmarks (Duffau, 2017; Ojemann et al., 1989). Thus, the neuronal organization of critical language-supporting areas should be functionally characterized at the individual level prior to removal of brain tissue affected by a disease such as epilepsy or tumor.

To aid neurosurgical decision making, functional speech cortical mapping (SCM) is usually performed invasively by electrical direct cortical stimulation (DCS) during awake surgery (Keles et al., 2004; Ojemann et al., 1989; Picht et al., 2006). DCS is currently the gold standard in cortical mapping, but it is limited by its invasiveness and spatial restriction to the immediate area around the surgical locus. Accurate noninvasive mapping would reduce time, costs, and risks in the operating room by guiding the DCS locations, and provide an alternative for patients who are not suitable for awake surgery.

To this end, transcranial magnetic stimulation (TMS) has been suggested as a noninvasive clinical tool to assess cortical areas critical to speech functions (Picht et al., 2013; Sollmann et al., 2015; Tarapore et al., 2013). Speech mapping with navigated repetitive TMS (nrTMS) is based on the virtual lesion approach (Silvanto & Cattaneo, 2017; Ziemann, 2009), where the nrTMS pulses are focused on the brain areas assumed relevant for speech processing while the patient performs an overt language task such as picture naming, action naming, or reading (Epstein et al., 1999; Hernandez-Pavon et al., 2014; Lioumis et al., 2012, 2023; Pascual-Leone et al., 1991). Applying a TMS pulse to speech-related cortical areas at the appropriate time can induce speech errors: these areas are thus considered relevant for the task (Krieg et al., 2017). Neuronavigational systems enable anatomically precise targeting of the TMS pulse (Hannula et al., 2005; Krings et al., 1997), and combining TMS with an audiovisual recording enables an accurate off-line evaluation of the subject’s performance (Lioumis et al., 2012). SCM by nrTMS is a clinically approved tool for presurgical evaluation of patients whose surgical locus lies near classical language areas especially in the left hemisphere (Jeltema et al., 2021; Mäkelä & Laakso, 2017), and for determining hemispheric lateralization of speech (Noorizadeh et al., 2024; Rezaie et al., 2020). Its objective is to provide a safe, reliable, and non-invasive *a priori* information on the arrangement of the cortical speech-related network. By providing such information to the surgical team, nrTMS-SCM can reduce the volume of craniotomy and the amount of postoperative language deficits (Sollmann et al., 2015).

Comparisons between DCS and TMS in SCM have shown that TMS-based speech mapping generally exhibits moderate-to-high negative predictive values (74.1%-100%) and sensitivity (68%-90.2%; see, for example (Krieg et al., 2014; Lehtinen et al., 2018; Picht et al., 2013; Tarapore et al., 2013)). SCM by TMS thus produces false positives but relatively few false negatives compared to DCS (Bährend et al., 2020; Picht et al., 2013; Tarapore et al., 2013). However, these comparisons remain limited, as DCS is typically confined to the cortical region immediately surrounding the resection. In addition to the large individual variance in the location of speech-related cortical sites, there is also significant variability in the temporal activation patterns across individuals (Levelt et al., 1998; Salmelin et al., 1994; Sörös et al., 2003). Individualizing the picture-to-TMS interval (PTI) to account for this variability could potentially increase the amount of evoked errors and the power of SCM by TMS. Despite its potential, the use of individualized PTIs in SCM TMS remains largely unexplored. Shinshi et al., 2015 reported significant increases in speech reaction times within PTIs between 150 and 375 ms when stimulating the left IFG, with 9 of 12 subjects having a significant increase in reaction time at 300 or 375 ms PTI. Sollmann et al., 2017, on the other hand, found a group-level decrease in error rates when increasing the PTI from 0 to 500 ms at 100-ms steps, but individual level results were not reported.

Accurate individualized PTIs require individual-level functional information from a modality with high temporal resolution, such as magnetoencephalography (MEG) or electroencephalography (EEG). Earlier M/EEG studies have demonstrated that speech production engages multiple brain regions with temporally overlapping activity; see, e.g., Ala-Salomäki et al., 2021; Liljeström et al., 2009; Vihla et al., 2006. Evoked MEG responses during picture naming have a good test-retest reliability, demonstrating their great utility for language brain network studies (Ala-Salomäki et al., 2021).

So far, only few studies have combined MEG and TMS in a speech mapping context.Tarapore et al., 2013 found locations of MEG oscillatory activity in the beta-range (15–30 Hz), indicative of speech motor processing, to concord well with TMS error locations in 5 out of 12 subjects. In contrast, Shinshi et al., 2015 found that, in the left IFG, the timing 40-Hz rTMS pulses that produced a significant slowing in speech reaction times was significantly correlated with the peak latency of the MEG event-related desynchronization in the 25–50 Hz frequency range.

Unlike previous studies that primarily examined spatial or temporal correspondence between MEG and TMS, in the present study, we investigated whether MEG data collected during picture naming could be used *a priori* to prospectively guide the timing of TMS stimulation. We hypothesized that a personalized approach—integrating both anatomical and functional information from each individual could improve the accuracy and impact of SCM nrTMS. To test this, we performed SCM with nrTMS at and around the left perisylvian cortex in 13 healthy subjects. Stimulation was applied using both common (0, 200, 300 and 400 ms) and individualized custom PTIs, the latter derived from MEG recordings during a picture naming task. We aimed to probe whether stimulation at individualized PTIs would induce higher error rates overall or at some specific cortical areas, and to inspect the spatiotemporal correspondence between MEG and SCM nrTMS. At the group level, we found a significant positive correlation between the latency of the MEG peak activation and the PTI that produced the highest error rate. Specifically, the MEG peak activation preceded the optimal PTI by an average of 132 ms in the whole stimulation area in the lateral left hemisphere and by 103 ms in the frontal left hemisphere. Additionally, we found 17 PTI and stimulation area combinations across nine subjects for which the error rate increased significantly compared to the average error rate of each subject. These findings suggest that optimal PTIs are highly individual, and mapping the peak latency of speech-related cortical activity with MEG—or potentially EEG—could serve as a practical guide for selecting optimal PTIs for each individual for TMS-based SCM in both clinical and research applications.

## Materials and Methods

### Subjects

13 healthy subjects participated in the study (age 25.8 ± 5.1 years (mean ± SD), range 21-40, 7 females) and provided their written informed consent. None of the subjects had neurological or psychiatric disorders, medications that affected the central nervous system, known cases of epilepsy in their family, or cardiac diseases. All subjects were right-handed, spoke Finnish as their native language and had normal or corrected-to-normal vision, and normal color vision. The measurement protocol was approved by the Helsinki University Hospital (HUS) Regional Committee on Medical Research Ethics.

### Experimental methods

The study consisted of three measurement sessions: anatomical MRI (magnetic resonance imaging), MEG with picture naming tasks, and nrTMS SCM with the same picture naming task (Fig. 1) as in MEG. Picture naming tasks in MEG and nrTMS SCM were conducted in Finnish. Data were collected successfully for all subjects in all measurements.

**Figure 1:**
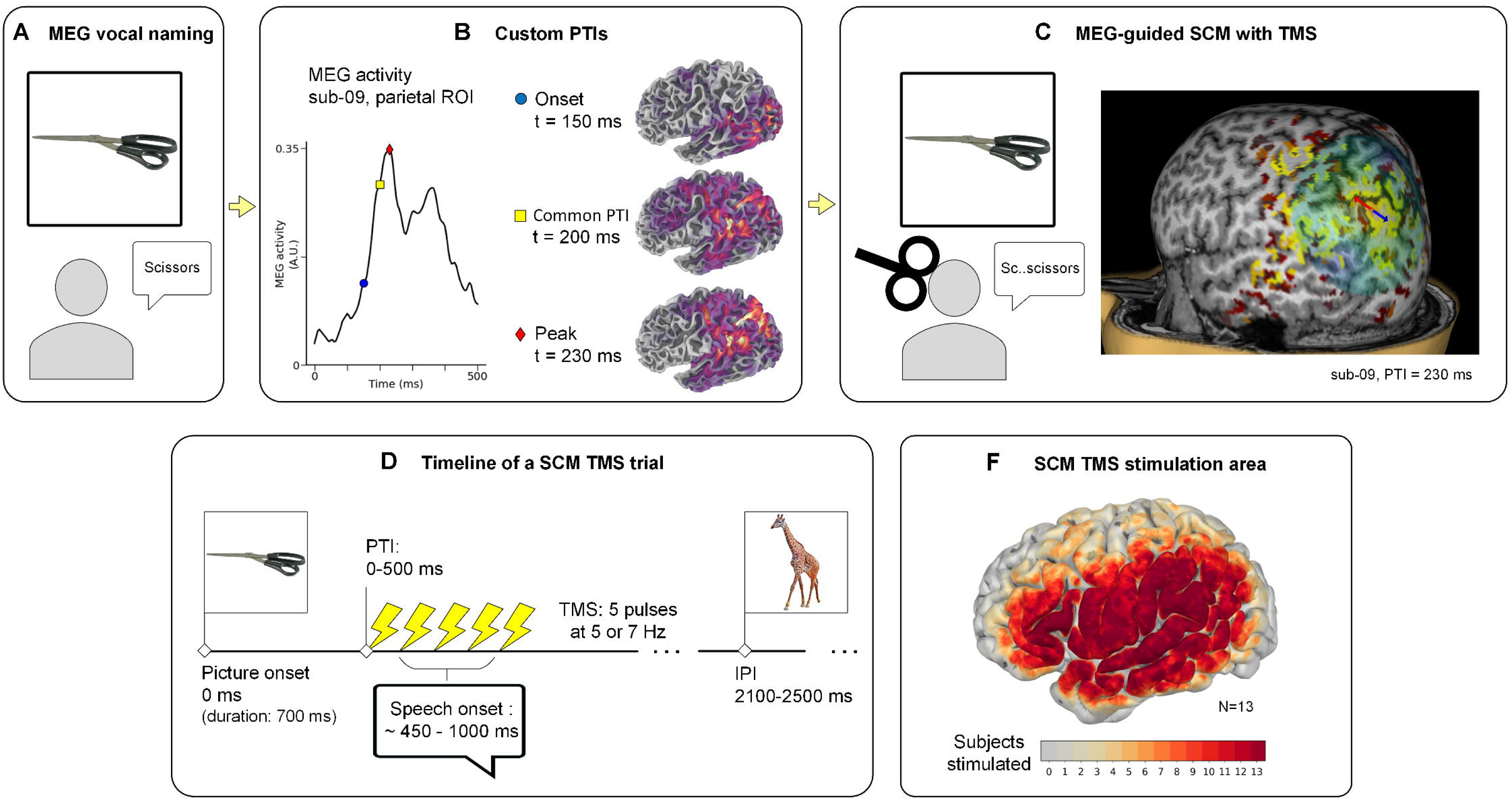
Study schematic. A) A vocal picture naming task was used to capture the cortical temporospatial patterns of speech production using magnetoencephalography (MEG). B) Customized picture-to-TMS (transcranial magnetic stimulation) intervals (PTIs) were chosen based on individual MEG data, in addition to common PTIs. Here, 200 ms is an example of a common PTI, and t=150 ms and t=230 ms are chosen customized PTI, corresponding to onset and peak of parietal region-of-interest (see Fig. 2 B for ROI) activity of this subject. C) Speech cortical mapping (SCM) was conducted with TMS (transcranial magnetic stimulation) using both common and customized PTIs. PTI-specific MEG data was overlaid on individual structural brain images for additional visual guidance. Here, thresholded MEG-data at 230 ms for sub-09 is overlaid. D) Timeline of one SCM trial. Each trial started with the presentation of a picture to be named, displayed for 700 ms. The PTI was changed between runs (7–9 runs per subject) and varied between 0 and 500 ms. The TMS pulse sequence consisted of 5 pulses, given at 5 Hz or 7 Hz stimulation frequency (inter-stimulus-interval of 200 ms or 143 ms). Speech onsets were measured from the throat with an accelerometer. The inter-picture-interval (IPI), or the time between consecutive trials, varied between 2100 and 2500 ms, depending on the subject and the run. F) Group level SCM TMS stimulation area. Coloring indicates the number of subjects who were given at least one TMS pulse around a cortical location during speech mapping (within 3 mm radius).

### MRI acquisition

MRIs (T1 and T2) were obtained for each subject. Four subjects were imaged in Helsinki University Hospital (HUS, Finland) using a 3T Siemens Skyra device (32-channel head-coil). Nine subjects were imaged in Aalto University’s Advanced Magnetic Imaging center (Espoo, Finland), with a similar 3T Siemens Skyra device (32-channel head coil). The images were processed using the Freesurfer software (Dale et al., 1999) to construct brain surface models for each subject.

### MEG measurements

MEG measurements were conducted in BioMag laboratory (HUS Diagnostic Center, Finland), with a whole-head MEG device (Elekta Neuromag TRIUX, MEGIN Ltd, Helsinki, Finland) consisting of 306 channels (102 magnetometers, 204 planar gradiometers). The recordings were conducted in a magnetically shielded room (Euroshield, Eura, Finland). The data were recorded with a sampling frequency of 1000 Hz, with an in-device bandpass filter of 0.03-330 Hz. The subject’s head position was continuously measured during the recording using five head position indicator (HPI) coils. Physiological artifacts were monitored using one electrode pair attached on the clavicles to measure the electocardiogram (ECG), two electrode pairs around the eyes to measure the electrooculogram (EOG), and four electrode pairs placed on top of different facial muscles (orbicularis oris superioris, orbicularis oris inferioris, zygomaticus, and digastric muscle) to measure muscle activity during speech.

Each MEG measurement consisted of two parts: a picture naming part and a facial gesture task. The MEG measurements and preprocessing steps are described in detail in (Tuomaala et al., 2025), where a method to reduce the amount of speech artifacts in MEG was introduced. The picture naming part included three different tasks: vocal naming, silent naming, and picture observation. In the vocal naming task, subjects were instructed to name aloud the object depicted in the image shown on a reflector screen placed 2 meters in front of them. In the silent naming task, subjects were instructed to name the object silently in their mind, without moving any facial muscles. In the picture observation task, the objective was to look at the shown images but not to name the objects. Occasionally, a red circle was presented overlaid on the image (control image), in which case the subject was instructed to say “now” aloud. In the facial gesture task, the subjects performed ten different facial gestures designed to activate various facial muscles. The information obtained from the facial gesture, silent naming, and observation tasks was applied to reduce the speech-related muscle artifacts in the vocal naming task; for the implementation, see (Tuomaala et al., 2025). In this study, the artifact-cleaned data from the vocal naming task were used.

The MEG experiment was conducted using a block design, where each task was repeated 18–19 times in a block (+ 2 control images in the observation task). In each repetition, an image was shown for 500 ms, after which there was a 2500 ms (in the naming tasks) or 1500 ms (observation task) response time, followed by a fixation cross for 1000 ms before showing the next image. The three task blocks were presented in pseudo-randomized order in three sets of six task blocks, resulting in six repetitions of each task block during the whole experiment. Two different block orders were used, so that 6 subjects were shown block order 1, and 7 subjects were shown block order 2. The block orders were chosen such that each of the three sets started with a different task. Including a short rest period between blocks, each measurement set (consisting of 6 blocks) was approximately 8 minutes long. The subjects had a short break between the measurement sets.

A set of 122 images was used in all naming and observation tasks during MEG. The images were normalized color images of living and non-living objects, from the bank of standardized stimuli (BOSS) (Brodeur et al., 2010, 2014). The chosen image set was optimized for native Finnish speakers, the images having high naming agreement and 1-4 syllables per word. In the MEG measurements, 110 images were all vocally named, silently named, and observed, with each image being presented once in each of the three different tasks. The remaining 12 images were control images for the observation task, and were not named. The same image set was later used during TMS speech cortical mapping.

Before the measurement, the subjects familiarized themselves with the images, and they were instructed to think of words that describe each object. We chose to let the subjects practice the images before the measurement to make the vocal naming process in MEG as similar as possible to the TMS experiment, where the subjects practice naming the images before any TMS pulses are given. The subjects were instructed to name the images at a brisk pace and to promptly move on to the next image if they were unable to correctly name a picture. Before the onset of the experiment, the subjects also trained the facial gestures, to ensure that each gesture was performed correctly. The subjects were monitored via a camera system during the MEG experiment and further instructed if tasks were not performed correctly.

### MEG preprocessing

The vocal naming data was preprocessed by first applying the temporally extended signal space separation (tSSS) algorithm (Taulu & Kajola, 2005; Taulu & Simola, 2006) to remove noise originating from outside the sensor array, and to transform each subject’s data to one common head position. FastICA (Hyvärinen, 1999) was used to remove cardiac and ocular artifacts using the measured ECG and EOG signals as references. The data was then downsampled to 200 Hz.

An automated algorithm was used to remove facial muscle artifacts from the MEG signals during vocal naming (see (Tuomaala et al., 2025) for details). The MEG and EMG signals were each decomposed using ICA, and mutual information (MI) between the ICA components of the EMG and MEG data was used to identify the muscle artifact components in the MEG data. After removal of artifact components, the cleaned MEG data was epoched and averaged.

MNE Python (Gramfort et al., 2013) was used for MEG source estimation. A boundary element model (BEM) was constructed from the Freesurfer surfaces to model the head conductivity, and used to define a source space with 20,484 sources (10,242 per hemisphere, ico5 spacing). MEG source reconstruction was performed using dynamical statistical parametric mapping (dSPM) (loose constraint = 0.3, depth weighting = 0.8). The noise covariance matrix was constructed using the baseline period (−0.2-0 s before each image onset) from the vocal naming epochs. Source reconstruction was performed on data from planar gradiometers. Subject-level data was further morphed to the fsaverage template brain surface for group-level visualizations.

We used each individual’s vocal naming MEG data to guide the planning of their TMS sessions, by selecting individualized picture-to-TMS intervals (PTIs) based on the timing of neural activity observed in MEG. MEG data was first inspected to yield an estimate of individual patterns of activation during the speech task, with the emphasis on finding time instants when activation within speech-related areas within the left hemisphere either emerged or peaked. Examples of customized PTI are shown in Fig. 1 B) and in Fig. 2 C). 3D MEG overlay-files were constructed for visualization to aid the TMS speech mapping (see Fig. 1 C)). This was done by filling a cortical ribbon mask generated during the Freesurfer surface generation with values corresponding to the closest MEG source strength (thresholded dSPM) at a chosen time.

**Figure 2:**
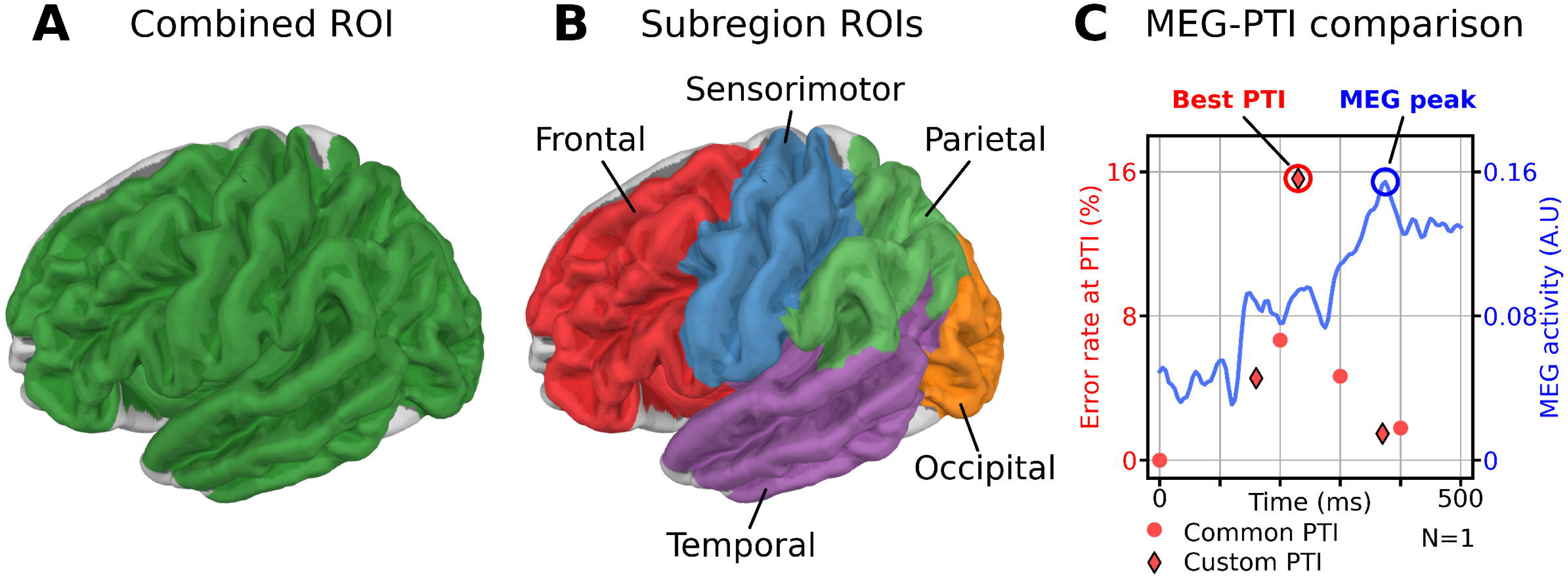
Regions of interest (ROIs) and MEG-PTI (magnetoencephalography, picture-to-TMS interval) comparison method. A) The combined ROI (CROI), which consists of all parcels that were targeted with TMS (transcranial magnetic stimulation) at least once. B) The five subregions forming the CROI were also inspected separately. C) Identifying the MEG response peak and the best PTI. For each subject and ROI, we estimated the MEG response peak and the best PTI using data only from that ROI. Here, data from frontal ROI for one exemplary subject is shown. Blue: the average MEG time course (normalized dynamical statistical parametric mapping values) at the frontal ROI, with the MEG peak activation circled. Red: the error rates for the different PTIs, including only the speech cortical mapping trials that targeted the frontal ROI, with the best PTI circled. Common PTIs (0, 200, 300, and 400 ms) are marked with red circles, and custom PTIs are marked with red diamonds and a black outline.

### TMS measurements

The TMS measurements were conducted in BioMag laboratory (HUS Diagnostic Center, Finland), using a navigated Nexstim NBS 5.2.4 system, with a figure-of-eight coil and an audiovisual recording of the subject’s performance. Each TMS session started with mapping the motor representation area of the right abductor pollicis brevis (APB), and defining a resting motor threshold (rMT). A stimulation intensity that evoked a potential of at least 50 *µ*V for 10 out of 20 pulses was chosen for rMT.

At the start of the session, the subjects were familiarized with the object naming image set, after which 2–3 baseline naming runs were conducted without TMS. During the baseline naming, any images that the subjects had trouble naming (incorrect word, stutter, delayed naming, anomia, etc.) were removed from the image set. The pacing of the images was adjusted during the baseline naming by changing the inter-picture-interval (IPI) so that the pace was slightly challenging for the subject. The IPI varied between 2100 and 2500 ms depending on the subject. In 9 out of 13 subjects we also attached an accelerometer (ADXL330 iMEMS Accelerometer, Analog Devices, Norwood, MA; 3000 Hz sampling frequency) on the throat either adjacent to or over the larynx to detect the speech onset.

During SCM TMS, a large area in the left hemisphere was stimulated, including the inferior frontal gyrus (IFG), supramarginal gyrus (SMG), angular gyrus (anG), superior and middle temporal gyrus (STG and MTG), and the mouth somatomotor area (see Fig. 1 E for group-level map of the stimulated areas). Some subjects were also stimulated in other nearby areas, such as the middle and superior frontal gyrus or the posterior parietal cortex, if their MEG data had shown activity in that area. A biphasic pulse sequence with 5 pulses at 5 or 7 Hz stimulation frequency was used in all SCM trials, with the first pulse of the sequence delivered at PTI. Each SCM session was started with the stimulation intensity of 100 % rMT.

The SCM TMS measurement sessions consisted of 7–9 runs with the PTI changing between the runs. Common PTIs of 0 ms, 200 ms, 300 ms, and 400 ms were used for all subjects. In addition to these, each subject was stimulated with 3–5 customized PTIs based on their MEG data. Custom PTI selection is visualized in Fig. 1 B), and custom and common PTIs with respect to MEG time series are shown in Fig. 2 C). The timeline of one SCM TMS trial is shown in Fig. 1 D. The used PTIs, IPIs, and stimulation frequencies with the corresponding speech error rates for each subject and each run are listed in Supplementary Table 2.

Some adverse effects were observed during SCM TMS. Based on the video analysis, seven subjects experienced visible pain, and eight subjects experienced visible muscle stimulation that affected their ability to speak during parts of the SCM exam. In three subjects, the stimulation intensity had to be lowered when stimulating the frontal lobe and the anterior temporal lobe to reduce pain (see Supplementary Table 2). In two subjects, the last measurement runs were visibly affected by tiredness, which resulted in inflated speech error rates in those runs (sub-04 and sub-08). When determining subject-specific runs with highest error rates, these runs were discarded and the second-highest error rates for these subjects were used instead. Tiredness also caused one of these subjects (sub-08) to miss one SCM TMS run altogether.

### Speech error analysis

The behavioral TMS data was processed using the recorded video and audio with an analyzer tool (NexSpeech Analyzer 2.1.0, Nexstim, Finland). Each naming trial was reviewed by a native Finnish speaker (S.A.) by comparing naming trials to the baseline naming trial, and then classified as a naming trial with error, naming trial with no error, or discarded. Naming errors were categorized to four categories: performance errors, no response errors, semantic errors, and other errors. The error categorization followed roughly the naming error categorization by (Corina et al., 2010).

During error review, we excluded trials with unsuccessful stimulation (coil movement, poor contact, or incorrect delivery). We also excluded naming-error trials affected by external factors (e.g., operator entering subject’s view), disturbing muscle stimulation or pain, or subject behaviors not related to naming (e.g., laughter). Additionally, we removed errors likely caused by carryover from the previous trial, and for repeated identical semantic errors (≥3), we kept only the first instance. Excluded trials were not included in any further analysis. Naming trials that were not classified as errors and were not discarded, were classified as non-errors. After classifying all naming trials, we merged the behavioral and stimulation data, linking each trial to the location and orientation of the stimulation coil and the estimated magnitude of the electric field (V/m) at each locus.

The accelerometer data was used to help identify the naming instances where the speech onset was delayed. The raw accelerometer data was first filtered with a 75-Hz high-pass filter to remove low frequency artifacts. Next, we computed the amplitude envelope of the rectified, filtered signal and applied a 5-sample moving average for further smoothing. From the smoothed envelope, speech onsets were found by locating the first sample where the signal value exceeded 5 *µ*V for at least 150 consecutive samples (50 ms with a sampling frequency of 3000 Hz). For each subject, we fit a log-normal to the speech-onset times and set the delay threshold to twice the distribution’s peak. The average threshold for a delay-type error was 1199 ms (N=9, range 1060-1390 ms), with a mean difference of 599 ms from the peak speech onset. In speech-error classification, delays were counted as performance errors.

### MEG and TMS analysis

To enable common analysis of TMS and MEG, the location and orientation of the TMS coil and the location of the TMS locus at individual peeling depth were transformed from the TMS system-defined voxel coordinates to Freesurfer-based MRI surface RAS coordinates used by MNE Python (Gramfort et al., 2013) to process source-reconstructed, surface-based MEG data. MNE Python was also used to visualize all surface-based brain images with MEG and TMS data, unless stated otherwise. The TMS locuses were projected on top of the gray matter surface along the axis between the coil location and the TMS locus in the brain volume. An example of the coordinate system transformation is shown in Supplementary Figure 1. TMS error counts, the error rate, and the error incidences (number of subjects with at least one error) were then calculated at the group level around each point on the gray matter surface within a radius of 10 mm. The error incidence was used to visualize the spatial distribution of errors at the group level at different PTIs/PTI ranges and for different speech error types. We chose to primarily use the error incidence in visualizing our group level data instead of error rate, since: 1) volume-based error rate calculation tends to produce inflated values on the edges of the stimulation area (since the number of SCM trials decreases at the edges of the stimulation area) and distort the results, and 2) the TMS error incidence heatmap visualizes TMS-positive sites across subjects and emphasizes subjects with low error counts, who are not well represented by group-level error count and error rate figures.

For spatiotemporal comparison of MEG and TMS, two approaches were adopted. First, cosine similarity was used to compare MEG activation at each time point with TMS error locations at the group level. Cosine similarity calculates the cosine of the angle between two vectors, with the result varying between ™1 (vectors having opposite directions) and 1 (vectors having the same direction), regardless of the vectors’ magnitudes. To place TMS and MEG in the same vector space, we mapped TMS error incidence onto the MEG source grid (10,242 locations on the left-hemispheric white matter surface). For each MEG vertex, we counted the TMS errors within a 10-mm radius on the gray matter surface. The error incidence at MEG source locations was calculated for all errors at the group level and separately for different PTIs/PTI ranges. The cosine similarity between the TMS and MEG data was then calculated for each time point of the MEG time series, to quantify how the cosine similarity changed over time with respect to the changing MEG activity. As the number of stimulated subjects differed by PTI, we matched the MEG average to the corresponding TMS sample size when needed.

Second, a region-of-interest (ROI) based approach was defined, using a custom anatomical parcellation by Aro, 2023. First, a ROI encompassing the whole stimulation (combined ROI, or CROI) was defined by including all parcels in the lateral left hemisphere that were targeted with TMS at least once (see Fig. 2 A). The parcels in the CROI were then divided into five subregions: frontal, sensorimotor, parietal, temporal, and occipital (see Fig. 2 B). The subregions roughly follow the anatomical lobes of the brain, with the exception of the sensorimotor region, which consists of the precentral gyrus and sulcus from the frontal lobe and the postcentral gyrus and sulcus from the parietal lobe. We defined five ROIs: four subregion ROIs (frontal, sensorimotor, parietal, temporal) and one combined ROI. The occipital subregion was included only in the combined ROI, as it was not systematically targeted by TMS and received thus only few stimuli.

For each ROI, the best PTI (PTI with the highest error rate) and the latency of the peak MEG response were individually determined. We then computed the Pearson correlation coefficient between these measures across ROIs. The best PTI per ROI was determined using only stimuli delivered over that ROI, and the peak MEG latency was derived from the ROI-averaged MEG time course. The procedure for determining the best PTI and peak MEG latency is illustrated in Fig. 2 C. Using the subregion ROIs, we tested with a one-tailed binomial test for each subject, whether stimulating over any subregion ROI at any PTI yielded significantly (p*<*0.05) more speech errors compared to the average error rate of the subject. Benjamini-Hochberg procedure was used to control for false discovery rate.

## Results

### MEG and TMS group-level results

Group-averaged MEG evoked responses in the vocal naming task peaked first bilaterally in the occipital cortex at ∼100-195 ms after the image onset, followed by activation in the middle-posterior temporal cortex and the parietal cortex, peaking at 200-295 ms (Fig. 3 A). Bilateral sensorimotor activation was first observed at ∼200-295 ms. After 300 ms, left-hemispheric temporal activation shifted anteriorly toward the mid-temporal region, accompanied by activation in the left IFG.

**Figure 3:**
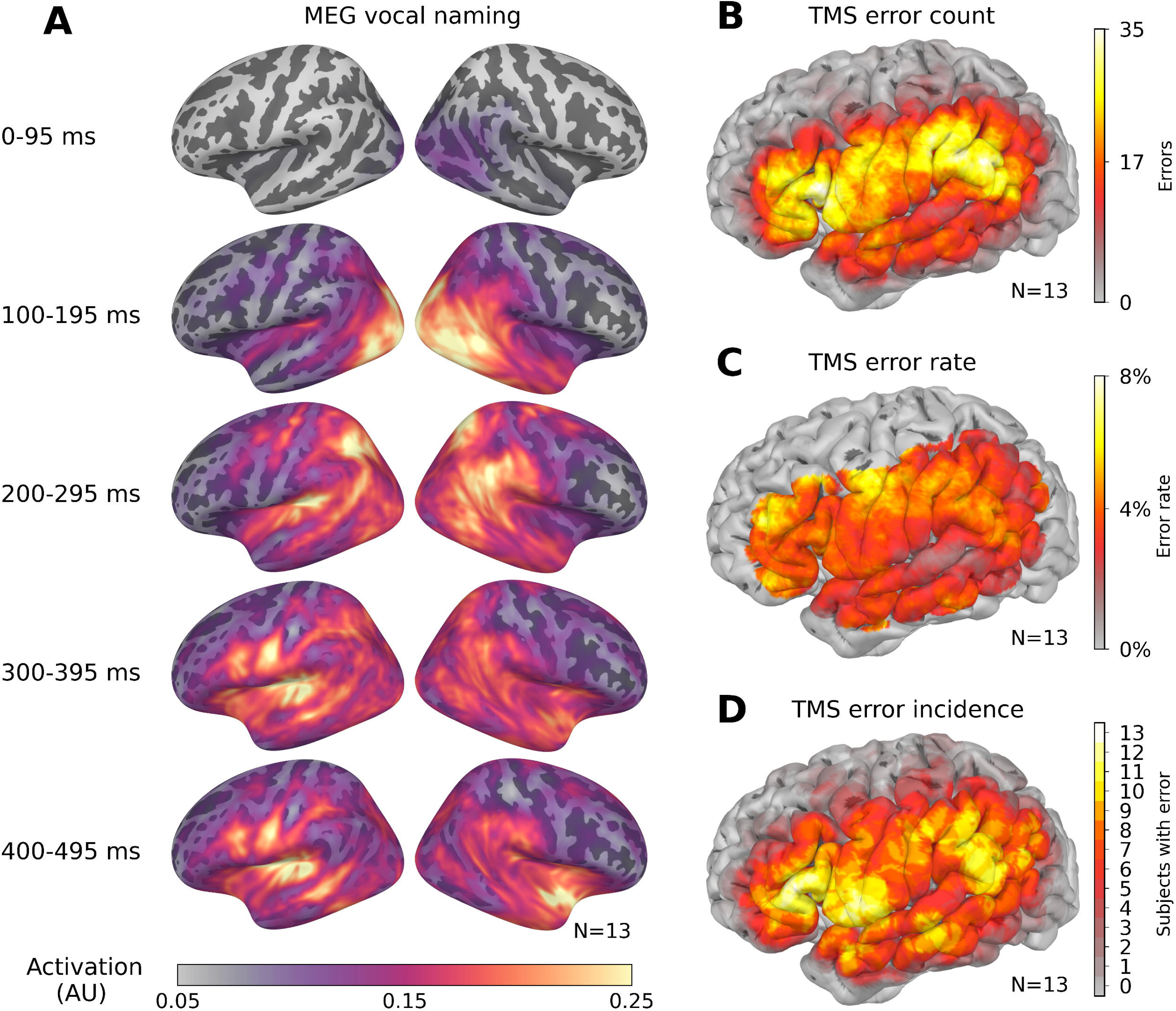
Group level (N=13) results from the MEG (magnetoencephalography) and TMS (transcranial magnetic stimulation) experiments. A) Group-average MEG evoked response in the left and the right hemisphere during vocal naming task (normalized dynamic statistical parameter mapping, dSPM). B) TMS error count, i.e., the number of errors observed per stimulation site. C) TMS error rate (N_errors_/N_stimuli_, [%]) per cortical location; locations with at least 200 stimuli given are visualized. D) TMS error incidence per cortical location, indicating how many subjects made at least one error. Number of errors and non-errors in B-D were counted within 10 mm radius from each cortical location.

On average, 1135 (range 893–1381) successful TMS pulse sequences were delivered during one measurement. All subjects made speech errors during the SCM. The average amount of speech errors was 39 (range 9–75), which corresponded to an average error rate of 3.6% (range 0.75–7.46%). Performance errors were the most common, consisting of 66.6% of all errors. No response and other errors were equally common, both representing 13.7% of all errors. Semantic errors were the rarest (6.1%). Supplementary Table 1 summarizes the number of speech errors per ROI and speech error type.

Fig. 3 B-D shows the group results from SCM TMS, including data from all runs and PTIs. Speech errors were found throughout the stimulation area, with the highest group level error count extending from the IFG to the sensorimotor cortex and SMG (see Fig. 3 B). The highest group-level error rates (see Fig. 3 C) were observed in the pars orbitalis, middle frontal gyrus, and precentral gyrus. However, since these areas are near the edge of the stimulation area, their values might be inflated due to edge-effects. Fig. 3 C) presents, for each cortical site, the number of participants who produced at least one speech error, identifying the sites most likely to elicit errors across participants. The most reliable sites were the pars triangularis area (13/13 subjects) and pars opercularis (12/13) which correspond to the classical Broca’s area. Stimulation in the inferior somatosensory and motor cortex near the orofacial representations evoked speech errors in 12/13 subjects. Another error hotspot was observed in the SMG and anG (11/13), near the classical Wernicke’s area. Finally, stimulation of the frontal and middle portions of STG and MTG elicited errors in 10/13 subjects.

### Effect of PTIs

Figure 4 shows, for each participant, the PTI (or PTI range) at which their highest error rate occurred, along with each participant’s mean total error rate (right) and the mean error rate for each PTI (or range; below). At the group level, speech errors were found at each PTI and PTI range, with the highest average error rates found in the 301-399 ms and 201-299 ms range. However, an independent two-sample permutation test (10,000 permutations) comparing mean group-level error rates found no significant differences between error rates across PTIs and PTI ranges (21 tests, smallest uncorrected *p*=0.07 (stat=0.0158) between 201-299 ms PTI range and 300 ms PTI).

**Figure 4:**
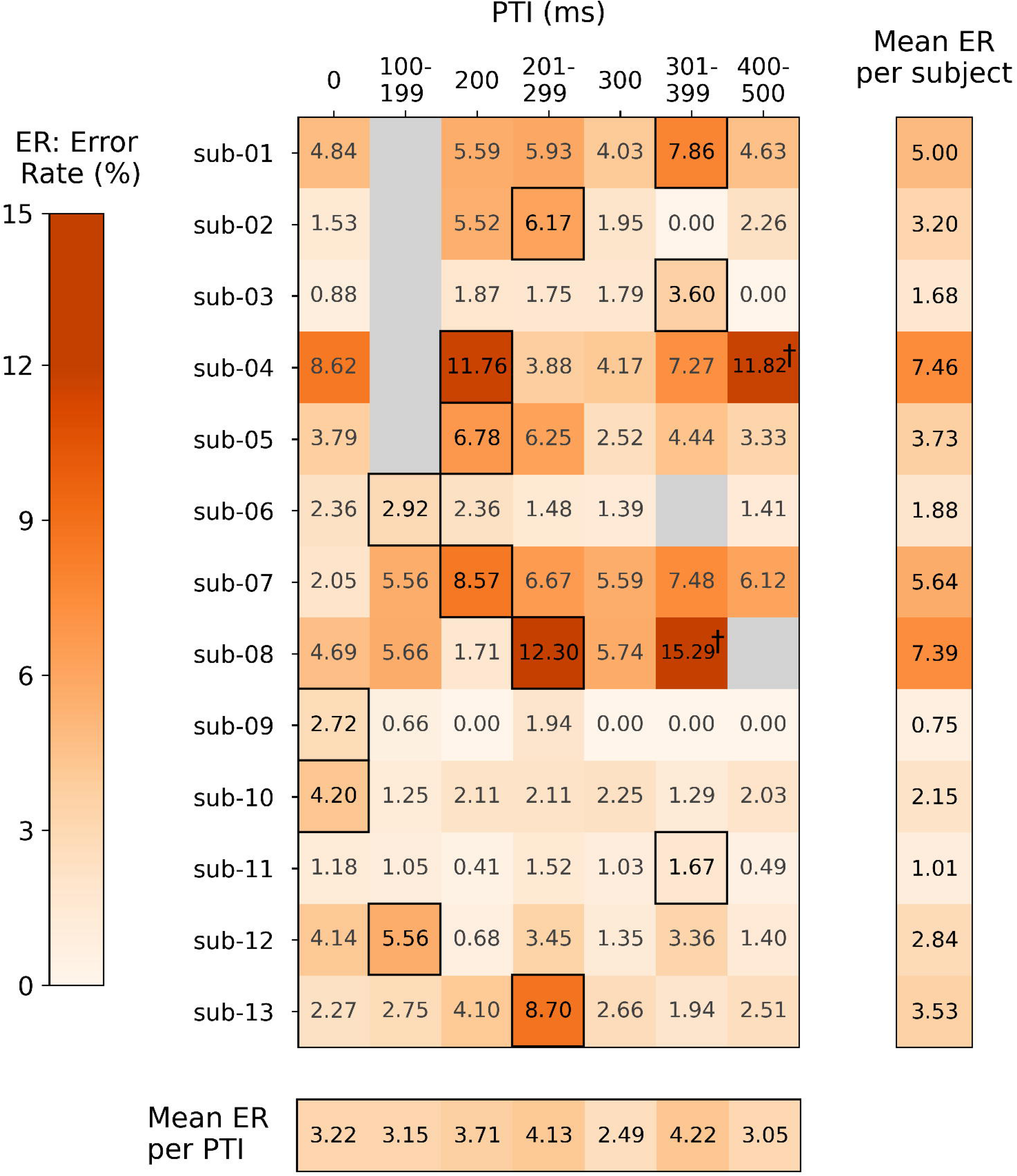
Speech error rates of each subject in each picture-to-TMS interval (PTI) range. Columns 0, 200, and 300 ms include errors from only that PTI, while columns 100–199, 201–299, 301–399, and 400–500 ms include the best error rate of the 1–3 custom PTIs in that time range. The best error rate PTI (or PTI range) of each subject is marked with a black box around the associated error rate value. Gray cells indicate missing conditions. Mean error rates of each subject (averaged over all PTIs) and mean error rates per PTI (averages over all subjects) are also visualized. †: Subject tired.

The PTI yielding the highest error rate varied widely across individuals: three participants had the peak at 200 ms, three in 201–299 ms, three in 301–399 ms, while two at 0 ms, and two in 100–199 ms. Notably, more than half of the participants (8/13) reached their maximum error rate at a custom PTI, i.e., using an individualized stimulation timing rather than a predefined bin. Four subjects had a relatively high average error rate (ER≥ 5%), four had a moderate (2.5% *>*ER*>* 5%) error rate, and in five subjects a low error rate (ER≤ 2.5%) was observed. However, a high error rate (ER≥ 5%) was observed for eight subjects at least once at some PTI.

Fig 5 shows the group-level error incidences at the PTIs and PTI ranges presented in Fig. 4, with the 201-299 ms PTI range split to roughly equalize the number of stimuli. While the exact spatial distribution of errors varied between the PTIs, stimulating the IFG and the SMG elicited speech errors at all PTIs in the group level. The spatial similarity of induced TMS errors across PTIs, based on the cosine similarity between group-level TMS error locations and group-level MEG activity in Fig. 5 I, was at its minimum at t=90 ms, during which only the unstimulated visual cortex was active. A local peak in cosine similarity is observed at t=210 ms, which corresponds to strong parietal activity and additional activation in the posterior temporal and sensorimotor regions. Finally, the global peak is achieved at t=340 ms, showing activation in the IFG, the temporal lobe, and the parietal lobe. Although the cosine similarity values vary in exact values for different PTIs, their temporal profiles have very similar shapes, quantitatively reflecting the spatial similarities seen in figures A-H. Other error hotspots were found with 300 ms PTI over the postcentral gyrus and SMG, and at 200 ms over the frontal sylvian fissure.

**Figure 5:**
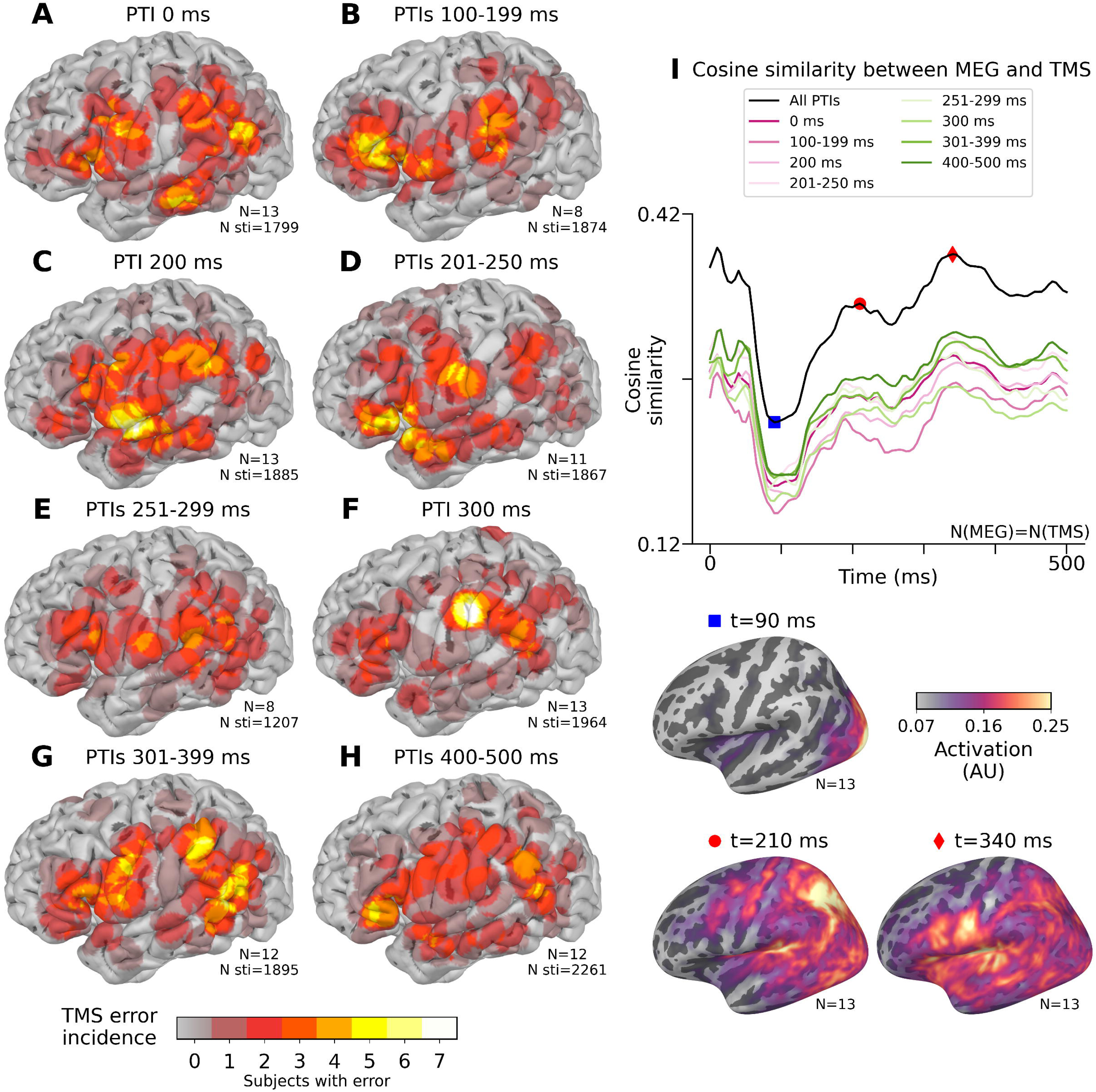
Group level TMS errors and cosine similarity compared to MEG at different PTIs. A-H: Group level TMS (transcranial magnetic stimulation) error incidence (number of subjects who made at least one speech error) at different PTIs (picture-to-TMS intervals). The number of errors and stimuli were calculated within a radius of 10 mm from each cortical location. I: cosine similarity between TMS errors and MEG activity. The cosine similarity was calculated for all TMS errors (at all PTIs, see fig. 3 D), and separately for all PTI ranges in (A-H). For each line, the MEG data included averaged data from only those subjects who were stimulated at that PTI or PTI range. Cosine similarity was calculated for each time point of the MEG time series. Global minimum of cosine similarity was reached at t=90 ms during occipital activation, and the local and global peaks were reached at t=210 ms and t=340 ms.

### MEG-TMS results

The comparisons of individual peak MEG latencies with the best TMS PTIs are visualized in Fig. 6. The peak MEG latency and the best PTI over the whole stimulation area showed a statistically significant correlation (Pearson R=0.713, *p*=0.006, 95% bias-corrected and accelerated bootstrap confidence interval [0.26, 0.88]), with the subject’s best PTI preceding their MEG peak (mean difference 132 ms (SD=77 ms) between best PTI and MEG peak, t(24)=3.33, *p*=0.003), see Fig. 6). The same analysis conducted at the subregion level revealed a significant correlation between the optimal PTIs and MEG peak latencies at the frontal ROI (R=0.673, *p*=0.012, CI [0.08, 0.94]), likewise the best PTI preceding the MEG peak (*x*=103 ms (SD=70 ms), t(24)=3.02, *p*=0.006). In the other subregions (sensorimotor, parietal, temporal) no significant correlations were observed, but, on average, the best PTIs did precede the MEG peaks. See Table 1 for the list of R-values for all ROIs and Table 2 for the list of peak MEG latencies, best PTIs, and the difference between MEG latency and PTI for all ROIs.

**Figure 6:**
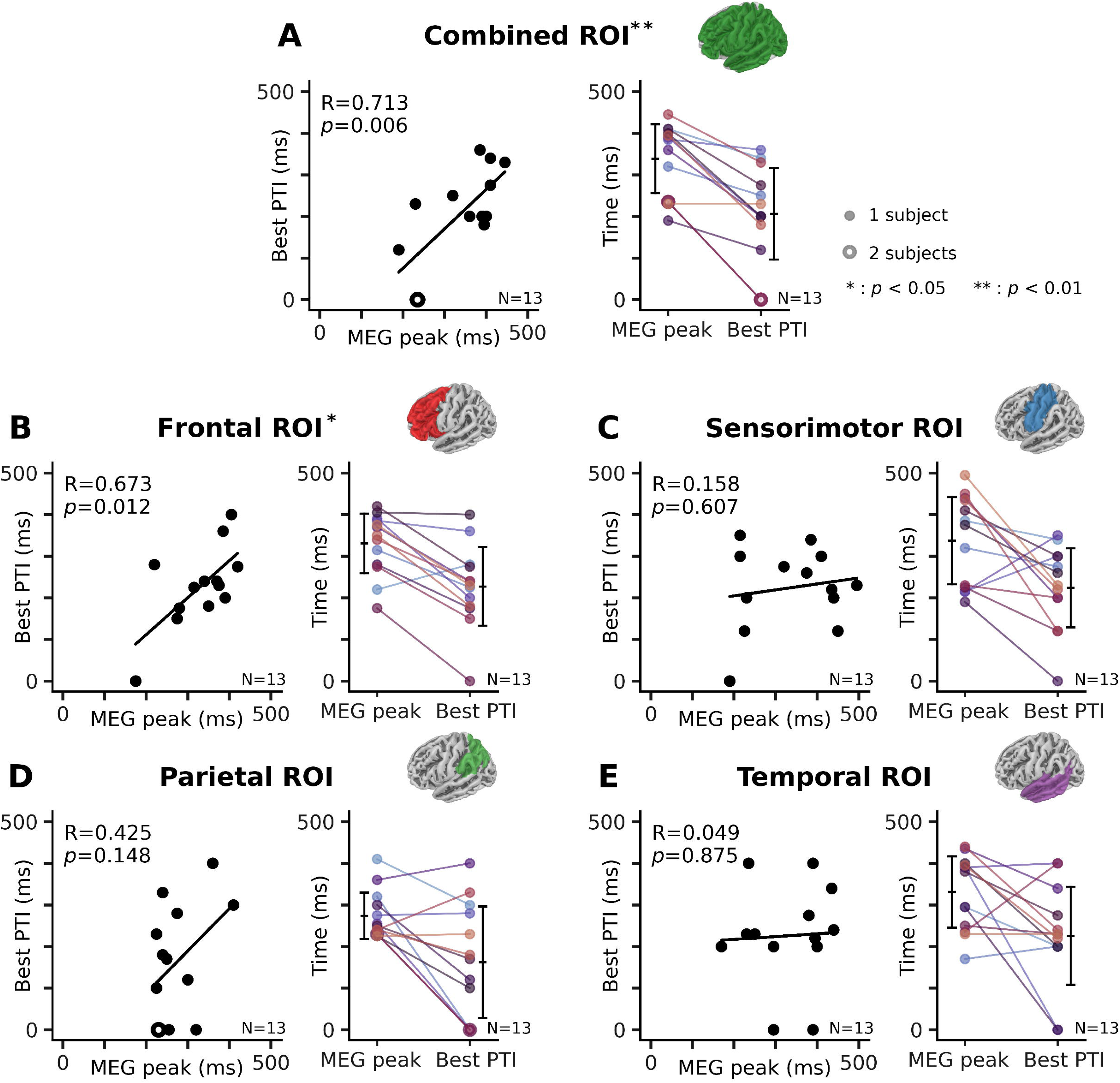
Comparisons of subject-specific peak time of MEG (magnetoencephalography) activity and the PTI (picture-to-TMS (transcranial magnetic stimulation) interval) with the highest error rate at different regions of interest (ROIs). The left panels of each subfigure show the line of regression and Pearson correlation coefficient (R) between the latency of MEG peak and best PTI, and the right panels show the means and the standard deviations of the latencies of MEG peak and best PTI. See also Table 1 for the list of R-values, and Table 2 for the list of means and standard deviations.

**Table 1:** Pearson correlation coefficients (R) with *p*-values and 95 % bias-corrected and accelerated bootstrap confidence intervals (CIs) between peak MEG (magnetoencephalography) latencies and best PTIs (picture-to-TMS intervals) in different regions-of-interest (ROIs) for N subjects. See Fig. 6 A-E left panels for the plots. CROI: combined ROI.

| ROI | R | $p$ | 95% CI | N |
| --- | --- | --- | --- | --- |
| CROI | 0.713 | 0.006 | [0.262, 0.883] | 13 |
| Frontal | 0.673 | 0.012 | [0.079, 0.940] | 13 |
| Sensorimotor | 0.158 | 0.607 | [-0.580, 0.720] | 13 |
| Parietal | 0.425 | 0.148 | [-0.362, 0.768] | 13 |
| Temporal | 0.049 | 0.875 | [-0.470, 0.490] | 13 |

**Table 2:** Comparisons of peak MEG (magnetoencephalography) latencies and best PTIs (picture-to-TMS intervals) in different ROIs (regions-of-interest). Average (*x*) and standard deviations (SD) for MEG peaks, best PTIs, and time difference between MEG peaks and best PTIs are given in milliseconds. Student’s t-test statistic (24 degrees of freedom) and the corresponding *p*-value for the difference in means of MEG peaks and best PTIs is reported. See Fig. 6 A-E right panels for the plots. N: number of subjects, CROI: combined ROI

| ROI | MEG peak (ms)<br>$\bar{x} \pm \text{SD}$ | Best PTI (ms)<br>$\bar{x} \pm \text{SD}$ | MEG peak - best PTI (ms)<br>$\bar{x} \pm \text{SD}$ | t(24) | $p$ | N |
| --- | --- | --- | --- | --- | --- | --- |
| CROI | 339 ($\pm 83$) | 207 ($\pm 110$) | 132 ($\pm 77$) | 3.327 | 0.003 | 13 |
| Frontal | 330 ($\pm 71$) | 227 ($\pm 95$) | 103 ($\pm 70$) | 3.025 | 0.006 | 13 |
| Sensorimotor | 337 ($\pm 105$) | 224 ($\pm 95$) | 113 ($\pm 130$) | 2.767 | 0.011 | 13 |
| Parietal | 274 ($\pm 56$) | 162 ($\pm 134$) | 112 ($\pm 121$) | 2.661 | 0.014 | 13 |
| Temporal | 331 ($\pm 86$) | 226 ($\pm 118$) | 105 ($\pm 142$) | 2.506 | 0.019 | 13 |

### Individual level results

Examples of individual level MEG and SCM TMS data are depicted in Fig. 7 for three subjects for the combined ROI. MEG and SCM TMS data are displayed at two time points: at the best PTI and at the MEG peak latency (also indicated in the MEG time series). The subjects exemplify the varying error rates observed in the study, with sub-06 having a generally low (1.88%), sub-04 having a high (7.46%), and sub-02 having a moderate (3.20%) error rate. They also have varying latencies for the best PTI and MEG peak, with sub-06 showing both the earliest latencies for MEG peak (190 ms) and best PTI (120 ms), and sub-04 showing a large time difference between the MEG peak and best PTI (190 ms difference, compared to 70 ms for the other subjects).

**Figure 7:**
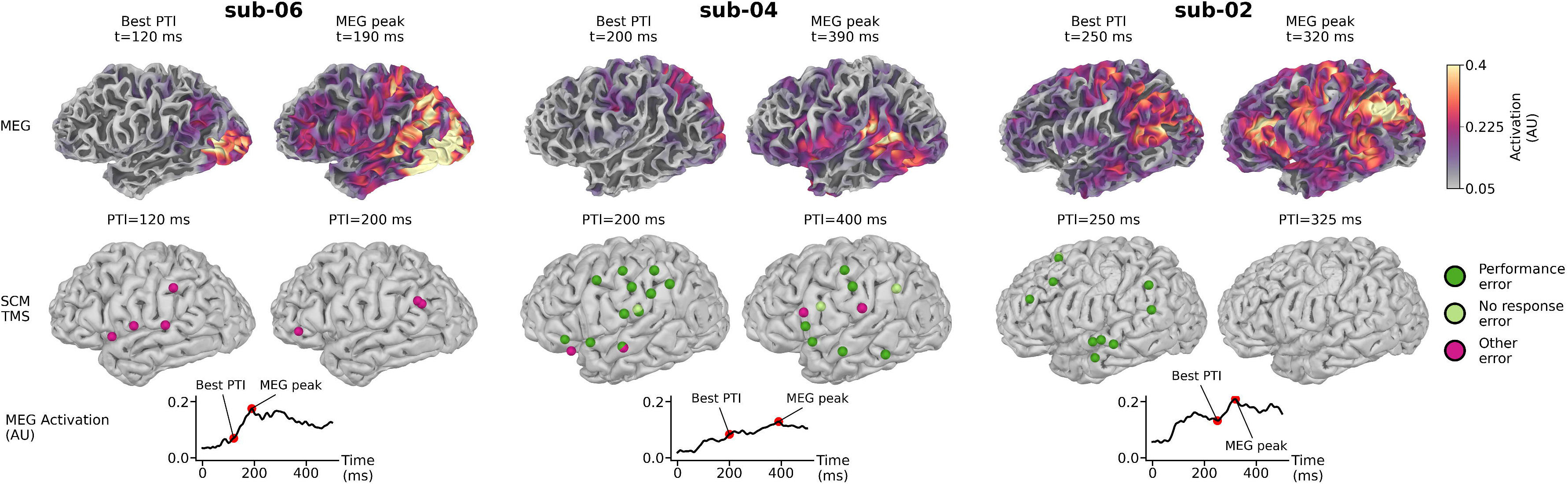
Individual-level MEG (magnetoencephalography) and SCM with TMS (speech cortical mapping with transcranial magnetic stimulation) results for three exemplar subjects. Top row: MEG source level data visualized on white matter surface. The left MEG image shows the snapshot of MEG activity at the time of the best PTI (picture-to-TMS interval), and the right shows the snapshot of MEG activity at the time of the MEG peak for each subject. The MEG peak indicates the peak of activity at the CROI (combined region-of-interest), which includes cortical areas in the left lateral hemisphere. Values are normalized dSPM values. Second row: SCM TMS data of one run using the indicated PTI. The left image shows the SCM TMS result of one run using the best PTI, and the right shows the SCM TMS result at a PTI nearest to the MEG peak in the CROI. SCM TMS error colors indicate the type of the speech error (dark green: performance error, light green: no response error, violet: other error). Third row: average of the MEG source time courses included in the CROI, with the best PTI and the MEG peak of each subject marked with a red dot.

Sub-06 shows strong MEG activation in the occipital lobe, and small activation in the SMG/anG at the time of best PTI (t=120 ms). 70 ms later at the MEG peak (t=190 ms), sub-06 shows strong occipital, parietal, and temporal activity, as well as some frontal and sensorimotor activity. Their SCM TMS speech errors are located around the sylvian fissure, extending from the SMG to the anterior sylvian point at PTI 120 ms, and at SMG/anG and pars triangularis at PTI 200 ms. Sub-04 shows small activity in the occipital, posterior parietal, and sensorimotor areas at the time of the best PTI (t=200 ms), while at the time of the MEG peak (t=390 ms) the activity is mainly present at the IFG, inferior sensorimotor area, and the posterior temporal lobe. The speech errors of sub-04 are present in the SMG, middle section of the postcentral gyrus, frontal TG, and near the IFG at both PTIs. For sub-02, the superior sensorimotor gyruses, posterior part of the STG, middle section of the TG, SMG, and anG activate at the best PTI (t=250 ms), while at the MEG peak (t=320 ms) the inferior and middle frontal gyruses and the inferior sensorimotor gyruses activate in addition to the previously activated areas. For the TMS data, sub-02 made speech errors when stimulated over the middle section of the temporal lobe, SMG, and the middle frontal gyrus at the best PTI (PTI=250 ms). Sub-02 did not make any speech errors with the PTI corresponding to the MEG peak (PTI=325 ms).

One-tailed binomial test revealed 17 significant subregion and PTI combinations, with at least one significant combination in nine of thirteen subjects (see table 3). Out of the 17 significant combinations, nine were located in the frontal ROI, four in the parietal ROI, three in the sensorimotor ROI, and one in the temporal ROI. The significant PTI’s occurred most frequently in the time window of 200–275 ms (n=8), followed by the 360–400 ms window (n=4), at 0 ms (n=3), and at 500 ms and 140 ms (n=1 each). Of these, one combination (sub-08, frontal ROI stimulation with 275 ms PTI) remained significant after correcting for multiple comparisons.

**Table 3:**
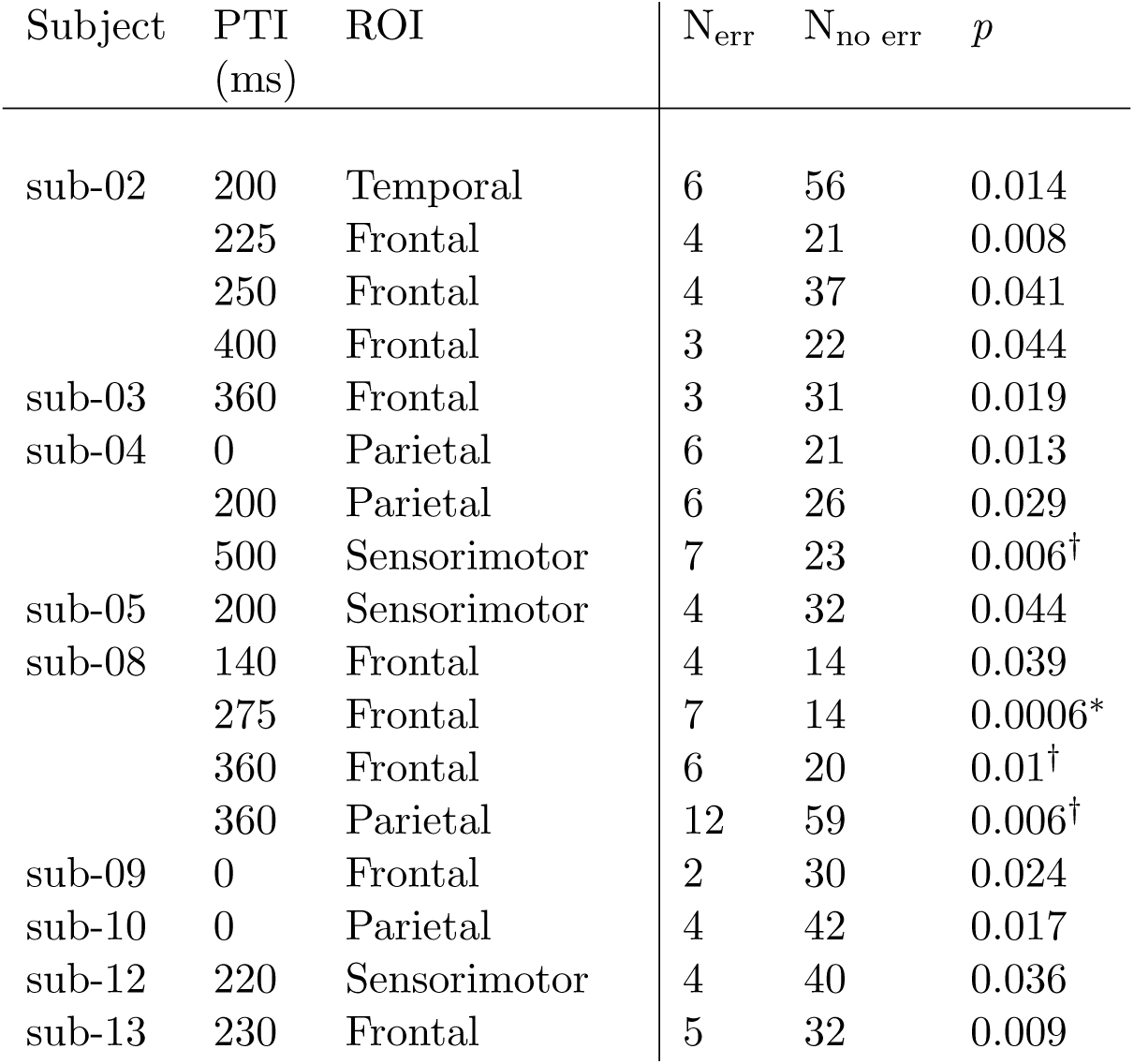
Results of one-tailed binomial tests (H_Alt_: *π>π*_0_) to find if stimulating over any cortical subregion ROI (frontal, sensorimotor, parietal, and temporal, see figure 2 B) at any PTI (picture-to-transcranial magnetic stimulation interval) yielded significantly (*p<*0.05) more speech errors compared to the average error rate of that subject. * : Significant after correcting for multiple comparisons (False discovery rate controlled with Benjamini–Hochberg procedure). † : Subject tired.

## Discussion

Our study demonstrates that magnetoencephalography can prospectively personalize the timing of navigated repetitive TMS (nrTMS) for speech cortical mapping (SCM). Across 13 healthy adults, the picture-to-TMS interval (PTI) that maximized TMS-induced speech errors correlated with each individual’s MEG peak latency, with the MEG peak preceding the optimal PTI on average by 132 ms across the lateral left hemisphere, and by 103 ms in the left frontal cortex. More than half of the participants reached their maximum error rate at a custom, MEG-derived PTI. At the group level, the error hotspots aligned with the canonical language sites. Together, our results support an MEG-informed, individualized approach to increase the sensitivity and utility of SCM nrTMS.

### MEG during overt picture naming reveals robust IFG activation after artifact suppression

Group-averaged MEG evoked responses during vocal picture naming showed the expected sequence of bilateral activation of the occipital, parietal, and temporal lobes, with additional left-hemispheric involvement of the IFG that appeared more pronounced in our study compared to previous findings(Liljeström et al., 2009; Vihla et al., 2006). Capturing IFG activation with M/EEG during overt speech is often hindered by the spatial overlap of cortical activation and speech-related myogenic artifacts in the frontal regions, motivating alternative mitigation strategies, such as silent (Liljeström et al., 2009) and delayed naming (Ala-Salomäki et al., 2021; Hultén et al., 2009). Despite using overt vocalization, our ICA and mutual information based artifact suppression (Tuomaala et al., 2025) seems to have efficiently attenuated the speech artifact, allowing the capturing of the frontal activation as well.

### Navigated rTMS reveals speech disruptions at the canonical cortical language areas

Group-level TMS errors spanned the entire stimulation area, and overlapped with the TMS-positive locations reported in earlier studies (Bährend et al., 2020; Krieg et al., 2014, 2016; Sollmann et al., 2015). The IFG, SMG, orofacial representation areas, and parts of TG were the most reliable sites to induce speech errors across participants, in line with the findings of Tarapore et al., 2013 who reported TMS-positive locations in the IFG, SMG, precentral gyrus, middle section of TG, and posterior STG.

### Effect of PTI and its individualization to SCM TMS

At the group level, the highest error rates were found at 201-299 ms and 301-399 ms PTIs, with no significant differences between the total error rates at different PTIs. This partly contradicts the results of Sollmann et al., 2017, who found 0 ms PTI to induce highest speech error rates overall. However, our observations on error subtypes converge: Sollmann et al., 2017 found only small differences in error rates across PTIs for hesitations and none for performance errors, aligning with our group-level data that consist mostly of delays and performance errors. Consistent with Shinshi et al., 2015, we also observed the highest error rates within later PTIs (≈300–400 ms).

Splitting the errors by PTI revealed that stimulating IFG and SMG elicited speech errors at all PTIs at the group level, suggesting that these regions are especially susceptible to interference across a wide temporal window. The spatial similarity of the error locations, confirmed by the similar shapes of cosine similarity between TMS at different PTIs and MEG, suggests that, despite heterogeneity in individual optimal timings, group-level interference maps align most closely with the canonical perisylvian language areas.

At the group level, the optimal PTI varied by ROI. For frontal, sensorimotor, and temporal ROIs the highest error rates were achieved with PTIs centered around 225 ms, while for parietal ROI the optimal PTI was on average ∼162 ms. This regional dissociation paralleled the MEG dynamics: the frontal, sensorimotor, and temporal ROIs peaked at ∼335 ms and the parietal region earlier at 274 ms. Notably, the mean difference between MEG peak and the best PTI was fairly constant across subregions, suggesting a stable delay between local neural activation and the most effective TMS interference window. These timings agree with earlier MEG reports of earlier parietal peak activations (280–300 ms; (Liljeström et al., 2009; Vihla et al., 2006)), followed by later temporal and frontal/sensorimotor engagement(∼370–500 ms; (Liljeström et al., 2009; Vihla et al., 2006)). The earlier best PTI for the parietal ROI followed by later best PTI for temporal and frontal ROIs suggests that the optimal PTIs reflect the underlying spatiotemporal progression of cortical activity and differ between cortical areas.

Our PTI findings may appear contradictory: At the group level, the locations of speech errors across PTIs look similar regardless of the chosen PTI. Yet, different cortical areas have different optimal PTI ranges. This is caused by the high interindividual variance in optimal timing: When data are averaged at fixed PTIs, participant-specific peaks occur at slightly different times and thus blur spatial differences, yielding similar incidence maps across PTIs. In contrast, when optimal PTIs are identified per participant and ROI, region-specific timing emerges. Therefore, group-level spatial stability across PTIs can coexist with ROI-specific, individualized optimal timings.

### Individualized PTIs improve sensitivity over fixed PTIs

Despite the absence of significant group-level differences in total error rates across PTIs, at the individual level 17 significant PTI-by-subregion combinations were observed. Significant combinations were found at all ROIs and they spanned a broad PTI time window (0-500 ms), with the frontal ROI (9/17) and the PTI time window of 200-275 ms (8/17) most commonly implicated. These findings confirm the importance of the IFG and PTIs around 200 ms as recommended sites and times for stimulation, but support the conclusion that different PTIs should be explored to find the PTI that is optimal for each individual and cortical area.

### MEG–PTI coupling: optimal stimulation precedes local MEG peaks

The observed correlations between best PTI and MEG peak latency, both across the whole measurement area (CROI) and in the frontal ROI, suggest an individual-level association between the time course of neuronal activity and the optimal time for TMS perturbation. These results are in concordance with those of Shinshi et al., 2015, who found a positive correlation between the MEG peak latency (event-related desynchronization, ERD) during picture naming and the TMS timing that significantly increased the reaction time in the left IFG. Our MEG peaks (330±71 ms) in the frontal ROI align well with their ERD peaks (typically 300-450 ms), but our best PTIs (227±95 ms) precede their reported optimal TMS timings (typically ∼300 ms). However, the studies are not entirely comparable since we examined not only the changes in reaction time but also other behavioral effects.

On average, the best PTIs preceded the corresponding peak MEG activity by about 100–130 ms. Several mechanisms can contribute to this finding. A local neuronal process disturbed by TMS at its onset or early phases could be directly impaired and prevented from resuming normally (the virtual lesion approach (Pascual-Leone, 1999)). Secondly, as communication between neuronal groups can be facilitated through neuronal coherence (Fries, 2005) and TMS can alter neuronal rhythms and phase alignment (Fuggetta et al., 2008; Thut et al., 2011), TMS could prevent downstream neurons from receiving information from regions that are active earlier. Such task-related oscillatory processes during speech processing have been observed earlier by, e.g., (Laaksonen et al., 2012; Liljeström et al., 2015; Piai et al., 2015). Furthermore, the perturbation caused by TMS can spread along structural neuronal networks (Esser et al., 2006) via white matter fibers. By this mechanism, TMS could also disturb cortical activation remotely, even if the stimulated site and the observed site were not directly connected (Siebner et al., 2009). For example, stimulation of the STG has been shown to modulate activity in the IFG as measured by functional MRI Vasileiadi et al., 2025; these areas are known to be connected via the arcuate fasciculus (AF) and the superior longitudinal fasciculus (SLF) (Friederici, 2011). Finally, since we used a five-pulse TMS sequence, we did not only disturb the speech processes at the onset of the pulse sequence (at the PTI), but at four more instances at intervals of 200 or 143 ms (5 or 7 Hz. This may increase the chance of interacting with critical speech processing stages if the stimulation train starts earlier. Alhough the optimal PTIs preceded the MEG peaks, the optimal PTI was not always the earliest (0 ms), indicating that the optimal PTI is dependent on the underlying cortical activity.

### Limitations and future directions

This study has some limitations that should be considered when interpreting the results and planning their clinical translation. PTI coverage was uneven across the naming task, and our results especially for the parietal ROI could be affected by the choice of PTIs. After 0 ms, the earliest PTIs were set at 200 ms for five subjects and at 100-199 ms for eight subjects; thus no subject had PTIs sampled in the]0, 100[time window, and 8/13 participants were missing the]0, 150[ms window. The ROIs that activate earlier in the speech process, such as the parietal ROI, could have been suboptimally sampled, potentially biasing the optimal PTIs upward. Conversely, the PTIs in the 200-300 ms range were sampled more densely than other PTIs, which can lead to finding the highest error rates in that time window more often.

Speech TMS on the frontal and temporal lobes is generally challenging, as subjects often report stimulation near these areas, i.e., near the face, as uncomfortable, painful or it causes muscle stimulation (Krieg et al., 2016; Lioumis et al., 2012; Tarapore et al., 2013). In our experiment, the stimulation intensity was decreased in these areas for three subjects who reported pain, and errors that were clearly caused by pain or disturbing muscle stimulation were excluded from analysis. There is still a residual risk that some of the speech errors are not caused by cortical stimulation but reflect peripheral effects.

The study participants were healthy young adults with no prior speech impairment, while the population participating in SCM TMS before a neurosurgery includes adults, elderly (Krieg et al., 2014), and children (Lehtinen et al., 2018), who might already suffer from a speech impairment or other cognitive difficulties (Krieg et al., 2014; Lehtinen et al., 2018; Picht et al., 2013). Thus some of the used stimulation parameter values, such as the IPIs (here 2100-2500 ms) or the PTIs (0-500 ms), might be too fast-paced and should be adjusted to fit each patient’s capabilities Lioumis et al., 2023. We recommend using MEG or EEG for patient-specific PTI estimation.

Using healthy participants also meant that we could not validate the TMS speech error locations against DCS-positive locations. Individualized MEG-guided selection of PTIs should next be tested in the clinical population preparing for neurosurgery, with the possibility for post-operative comparison to DCS mapping. The higher error rates in SCM TMS do not always directly translate to more informative SCM, if e.g., the spatial distribution of the errors is large and the errors show little clustering or reproducibility. Krieg et al., 2014 compared SCM TMS with DCS, and found better SCM–DCS compatibility at earlier PTIs, particularly in frontal regions, which our results also support.

The number of participants in this study was relatively small (N=13), which contributed to the large confidence intervals in the obtained correlation coefficients (e.g. [0.26, 0.88] 95%-CI for the R=0.713 in the CROI). While the number of participants was in line with previous SCM TMS studies, see e.g., (Nettekoven et al., 2021; Shinshi et al., 2015; Sollmann et al., 2017) larger cohorts in future studies will enable more reliable subject-level inferences.

Based on our results, the optimal individual PTIs could in the future be predicted with a rather straightforward method by calculating the peak MEG timings from large surface-based ROIs. Such large ROIs spatially smooth both SCM TMS and MEG data, which–while increasing the signal-to-noise ratios of both–can lead to source mixing in MEG and clustering of far-off speech errors in SCM TMS. Finer parcellations might increase specificity but would require more stimuli per parcel to maintain statistical power. Additionally, larger ROIs also facilitate the use of EEG for which volume conduction further blurs spatial specificity (Hämäläinen et al., 1993). In the future, real-time EEG with the possibility to measure event-related potentials (ERPs) during the SCM could enable brain-state TMS (Ahola et al., 2025) and constitute a major paradigm shift towards brain-computer interface-based mapping (Matsuda et al., 2026).

To conclude, stimulating with individually chosen PTIs can increase the number of speech errors elicited during SCM TMS, and MEG can provide a practical guide for choosing those timings. Several PTIs should be explored to account for the large individual variance and the differences between the optimal stimulation time windows between different parts of the eloquent cortices.

## Acknowledgements

We thank Henri Lehtinen, Elena Ukharova, Juha Montonen, Ville Mäntynen, and Jari Kainulainen for help during the measurement and analysis process. We are very grateful to the volunteers who participated in our study.

The study was funded by HUS VTR (TYH2022224); Research Council of Finland (357660 to P.L. and S.V., UAK780VAAL, 355409 to H.R.); the Flagship of Advanced Mathematics for Sensing, Imaging, and Modeling (359181 to H.R.); the Sohlberg Foundation (to H.R.); Maire Taposen säätiö (to P.L.); the Sigrid Jusélius Foundation (to J.G. and H.R.); the Emil Aaltonen Foundation (to S.A. and J.G.).

