## Supplements for "MEG-informed navigated TMS for individualized speech cortical mapping"

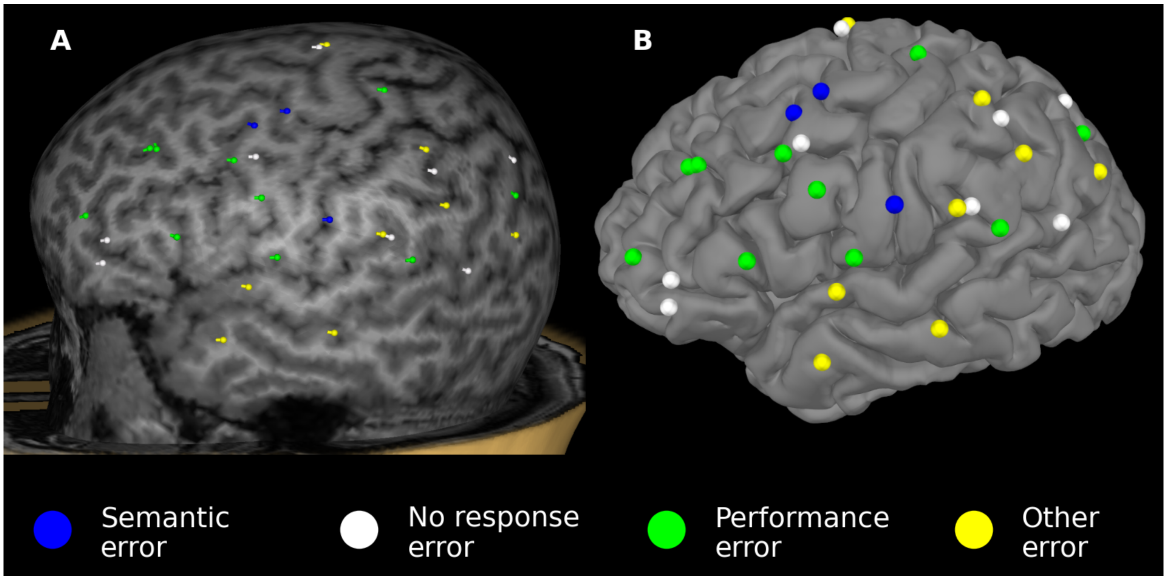


Supplementary Figure 1: Naming errors of subject 12 visualized in two coordinate systems, A) the original Nexstim coordinate system at peeling depth 21 mm (visualized in Nexstim NBS 5.2.4) and B) the Freesufer MRI surface RAS coordinate system, (visualized with MNE Python 1.6), with the TMS error locations projected on top of the gray matter surface. The errors in A) and B) are colored by the error category, using the default Nexstim color scheme.

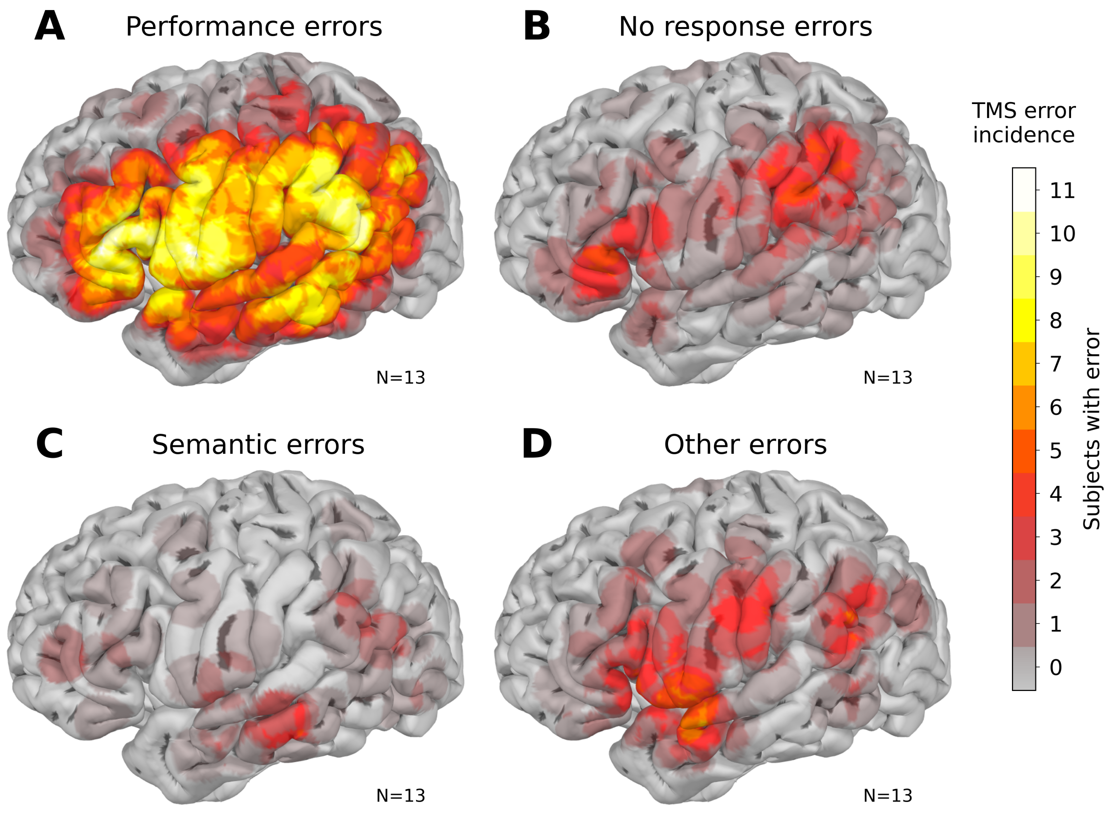


Supplementary Figure 2: Group level (N=13) error incidence by error type. Error incidence depicts how many subjects made at least one speech error of the specified error type within 10 mm radius from each cortical location.

| | Frontal | Sensorimotor | Parietal | Temporal | Occipital | Σ | % |
|---|---|---|---|---|---|---|---|
| Performance | 78 | 80 | 99 | 80 | 4 | 341 | 66.6 |
| No response | 20 | 15 | 23 | 11 | 1 | 70 | 13.7 |
| Semantic | 6 | 4 | 7 | 12 | 2 | 31 | 6.1 |
| Other | 15 | 19 | 21 | 13 | 2 | 70 | 13.7 |
| Σ | 119 | 118 | 150 | 116 | 9 | 512 | |
| % | 23.2 | 23.0 | 29.3 | 22.7 | 1.8 | | 100 |

Supplementary Table 1: Group level (N=13) error counts by error type and anatomical subregion.

| Subject | PTI (ms) | N err | N sti | ER (%) | IPI (ms) | SF (Hz) | SI (%) |
|---|---|---|---|---|---|---|---|
| **sub-01** | 0 | 6 | 124 | 4.84 | 2300 | 5 | 35 |
| | 200 | 8 | 143 | 5.59 | 2300 | 5 | 35 |
| | 260 | 7 | 134 | 5.22 | 2300 | 5 | 35 |
| | 280 | 8 | 135 | 5.93 | 2300 | 5 | 35 |
| | 300 | 5 | 124 | 4.03 | 2500 | 5 | 35 |
| | 340 | 11 | 140 | 7.86 | 2300 | 5 | 35 |
| | 400 | 12 | 259 | 4.63 | 2400 | 5 | 35 |
| | 440 | 3 | 141 | 2.13 | 2300 | 5 | 35 |
| (Σ, Σ, x̄) | | 60 | 1200 | 5 | | | |
| **sub-02** | 0 | 2 | 131 | 1.53 | 2200 | 5 | 33 |
| | 200 | 8 | 145 | 5.52 | 2200 | 5 | 33 |
| | 225 | 6 | 139 | 4.32 | 2200 | 5 | 33 |
| | 250 | 10 | 162 | 6.17 | 2200 | 5 | 33 |
| | 275 | 5 | 135 | 3.7 | 2200 | 5 | 33 |
| | 300 | 4 | 205 | 1.95 | 2200 | 5 | 33 |
| | 325 | 0 | 139 | 0 | 2200 | 5 | 33 |
| | 400 | 3 | 133 | 2.26 | 2200 | 5 | 33 |
| (Σ, Σ, x̄) | | 38 | 1189 | 3.2 | | | |
| **sub-03** | 0 | 1 | 114 | 0.88 | 2500 | 5 | 40 |
| | 200 | 2 | 107 | 1.87 | 2500 | 5 | 40 |
| | 240 | 4 | 229 | 1.75 | 2500 | 7 | 40 |
| | 280 | 2 | 115 | 1.74 | 2500 | 5 | 40 |
| | 300 | 2 | 112 | 1.79 | 2500 | 5 | 40 |
| | 360 | 4 | 111 | 3.6 | 2500 | 5 | 40 |
| | 400 | 0 | 105 | 0 | 2500 | 5 | 40 |
| (Σ, Σ, x̄) | | 15 | 893 | 1.68 | | | |
| **sub-04** | 0 | 10 | 116 | 8.62 | 2500 | 5 | 35 |
| | 200 | 14 | 119 | 11.76 | 2500 | 5 | 35 |
| | 220 | 4 | 105 | 3.81 | 2500 | 5 | 35 |
| | 260 | 5 | 129 | 3.88 | 2500 | 5 | 35 |
| | 300 | 4 | 96 | 4.17 | 2500 | 5 | 35 |
| | 350 | 8 | 110 | 7.27 | 2500 | 5 | 35 |
| | 400 | 10 | 126 | 7.94 | 2500 | 5 | 35 |
| | 500 | 13 | 110 | 11.82 | 2500 | 5 | 35 |
| (Σ, Σ, x̄) | | 68 | 911 | 7.46 | | | |
| **sub-05** | 0 | 5 | 132 | 3.79 | 2100 | 5 | 36 |
| | 200 | 8 | 118 | 6.78 | 2100 | 5 | 36 |
| | 240 | 8 | 128 | 6.25 | 2100 | 5 | 36 |
| | 280 | 5 | 129 | 3.88 | 2100 | 5 | 36 |
| | 300 | 4 | 159 | 2.52 | 2300 | 5 | 36 |
| | 340 | 6 | 135 | 4.44 | 2100 | 5 | 36 |
| | 380 | 2 | 125 | 1.6 | 2100 | 5 | 36 |
| | 400 | 4 | 120 | 3.33 | 2100 | 5 | 36 |
| | 440 | 2 | 134 | 1.49 | 2100 | 5 | 36 |
| (Σ, Σ, x̄) | | 44 | 1180 | 3.73 | | | |
| **sub-06** | 0 | 3 | 127 | 2.36 | 2200 | 5 | 33; 29 |
| | 120 | 4 | 137 | 2.92 | 2200 | 5 | 33; 29 |
| | 175 | 4 | 149 | 2.68 | 2200 | 5 | 33; 29 |
| | 200 | 3 | 127 | 2.36 | 2400 | 5 | 33; 29 |
| | 235 | 2 | 135 | 1.48 | 2400 | 5 | 33; 29 |
| | 280 | 1 | 158 | 0.63 | 2200 | 5 | 33; 29 |
| | 300 | 2 | 144 | 1.39 | 2300 | 5 | 33; 29 |
| | 400 | 2 | 142 | 1.41 | 2200 | 5 | 33; 29 |
| (Σ, Σ, x̄) | | 21 | 1119 | 1.88 | | | |

| Subject | PTI (ms) | N err | N sti | ER (%) | IPI (ms) | SF (Hz) | SI (%) |
|---|---|---|---|---|---|---|---|
| **sub-07** | 0 | 3 | 146 | 2.05 | 2300 | 5 | 33 |
| | 100 | 9 | 162 | 5.56 | 2300 | 5 | 33 |
| | 170 | 8 | 148 | 5.41 | 2300 | 5 | 33 |
| | 200 | 12 | 140 | 8.57 | 2400 | 5 | 33 |
| | 225 | 5 | 146 | 3.42 | 2300 | 5 | 33 |
| | 260 | 10 | 150 | 6.67 | 2300 | 5 | 33 |
| | 300 | 8 | 143 | 5.59 | 2300 | 5 | 33 |
| | 350 | 11 | 147 | 7.48 | 2300 | 5 | 33 |
| | 400 | 9 | 147 | 6.12 | 2300 | 5 | 33 |
| (Σ, Σ, x̄) | | 75 | 1329 | 5.64 | | | |
| **sub-08** | 0 | 6 | 128 | 4.69 | 2400 | 5 | 33; 29 |
| | 140 | 6 | 130 | 4.62 | 2400 | 5 | 33; 27 |
| | 170 | 9 | 159 | 5.66 | 2400 | 5 | 33; 29 |
| | 200 | 2 | 117 | 1.71 | 2500 | 5 | 33; 31 |
| | 275 | 15 | 122 | 12.3 | 2400 | 5 | 33; 29 |
| | 300 | 7 | 122 | 5.74 | 2400 | 5 | 33; 29 |
| | 360 | 24 | 157 | 15.29 | 2400 | 5 | 33; 29 |
| (Σ, Σ, x̄) | | 69 | 935 | 7.38 | | | |
| **sub-09** | 0 | 4 | 147 | 2.72 | 2300 | 5 | 34 |
| | 120 | 1 | 152 | 0.66 | 2300 | 5 | 34; 32 |
| | 150 | 1 | 162 | 0.62 | 2300 | 5 | 34; 32 |
| | 200 | 0 | 141 | 0 | 2300 | 5 | 34; 32 |
| | 230 | 3 | 155 | 1.94 | 2300 | 5 | 34 |
| | 300 | 0 | 151 | 0 | 2300 | 5 | 34; 32 |
| | 325 | 0 | 141 | 0 | 2300 | 5 | 34; 32 |
| | 400 | 0 | 149 | 0 | 2300 | 5 | 34; 32 |
| (Σ, Σ, x̄) | | 9 | 1198 | 0.75 | | | |
| **sub-10** | 0 | 6 | 146 | 4.11 | 2300 | 5 | 35 |
| | 150 | 2 | 167 | 1.2 | 2300 | 5 | 35 |
| | 200 | 2 | 144 | 1.39 | 2300 | 5 | 35 |
| | 240 | 2 | 145 | 1.38 | 2300 | 5 | 35 |
| | 300 | 4 | 181 | 2.21 | 2300 | 5 | 35 |
| | 320 | 1 | 158 | 0.63 | 2300 | 5 | 35 |
| | 400 | 3 | 149 | 2.01 | 2300 | 5 | 35 |
| (Σ, Σ, x̄) | | 20 | 1090 | 1.83 | | | |
| **sub-11** | 0 | 2 | 167 | 1.2 | 2200 | 5 | 35 |
| | 120 | 2 | 186 | 1.08 | 2200 | 5 | 35 |
| | 200 | 1 | 238 | 0.42 | 2200 | 5 | 35 |
| | 240 | 3 | 196 | 1.53 | 2200 | 5 | 35 |
| | 300 | 2 | 190 | 1.05 | 2200 | 5 | 35 |
| | 330 | 3 | 177 | 1.69 | 2200 | 5 | 35 |
| | 400 | 1 | 203 | 0.49 | 2200 | 5 | 35 |
| (Σ, Σ, x̄) | | 14 | 1357 | 1.03 | | | |
| **sub-12** | 0 | 6 | 145 | 4.14 | 2500 | 5 | 36 |
| | 180 | 8 | 144 | 5.56 | 2500 | 5 | 36 |
| | 200 | 1 | 147 | 0.68 | 2500 | 5 | 36 |
| | 220 | 5 | 145 | 3.45 | 2500 | 5 | 36 |
| | 300 | 2 | 148 | 1.35 | 2500 | 5 | 36 |
| | 340 | 5 | 149 | 3.36 | 2500 | 5 | 36 |
| | 400 | 2 | 143 | 1.4 | 2500 | 5 | 36 |
| (Σ, Σ, x̄) | | 29 | 1021 | 2.84 | | | |
| **sub-13** | 0 | 4 | 176 | 2.27 | 2500 | 5 | 31 |
| | 160 | 5 | 182 | 2.75 | 2500 | 5 | 31 |

| | | | | | | |
|---|---|---|---|---|---|---|
| 200 | 8 | 195 | 4.1 | 2500 | 5 | 31 |
| 230 | 16 | 184 | 8.7 | 2500 | 5 | 31 |
| 300 | 5 | 188 | 2.66 | 2500 | 5 | 31 |
| 370 | 4 | 206 | 1.94 | 2500 | 5 | 31 |
| 400 | 5 | 199 | 2.51 | 2500 | 5 | 31 |
| ($\Sigma$, $\Sigma$, $\bar{x}$) | 47 | 1330 | 3.53 | | | |

PTI: picture-to-TMS interval (ms)
ER: Error rate (%)
SI: stimulation intensity (% of max. stimulator output (max; min) )

N err: Number of errors
IPI: inter-picture-interval (ms)

N sti: Number of successfull stimuli
SF: Stimulation frequency (Hz)

Supplementary Table 2: TMS parameters of each subject.
